# Long-term single-molecule tracking in living cells using weak-affinity protein labeling

**DOI:** 10.1101/2024.07.11.603077

**Authors:** Claudia Catapano, Marina S. Dietz, Julian Kompa, Soohyen Jang, Petra Freund, Kai Johnsson, Mike Heilemann

**Affiliations:** Institute of Physical and Theoretical Chemistry, Goethe-University Frankfurt, Frankfurt, Germany; Department of Chemical Biology, Max Planck Institute for Medical Research, Heidelberg, Germany; IMPRS on Cellular Biophysics, Frankfurt am Main, Germany

**Keywords:** single-molecule fluorescence, single-particle tracking, self-labeling protein tags, weak-affinity binders, membrane receptors

## Abstract

Single-particle tracking (SPT) has become a powerful tool to monitor the dynamics of membrane proteins in living cells. However, permanent labeling strategies for SPT suffer from photobleaching as a major limitation, restricting observation times, and obstructing the study of long-term cellular processes within single living cells. Here, we use exchangeable HaloTag Ligands (xHTLs) as an easy-to-apply labeling approach for live-cell SPT and demonstrate extended observation times of individual live cells of up to 30 minutes. Using the xHTL/HT7 labeling system, we measure the ligand-induced activation kinetics of the epidermal growth factor receptor (EGFR) in single living cells. Furthermore, we generate spatial maps of receptor diffusion in cells, report non-uniform distributions of receptor activation, and the formation of spatially confined ‘hot spots’ of EGFR activation. This approach represents a general strategy to monitor protein dynamics in a functional context and for extended observation times in single living cells.

## Introduction

The plasma membrane of a cell constitutes a dynamic interface between the intra- and extracellular space and serves as a gate for information exchange. Embedded within this lipid bilayer are membrane receptors, the key elements of this communication, that respond to ligand binding by forming multi-protein complexes and initiate a signaling cascade inside the cell.^[1,2]^ Membrane receptor activation involves intricate interactions with other membrane proteins and lipids. Thus, functional characterization requires methods that allow the visualization of living cells.

Single-particle tracking (SPT) has developed into a powerful tool to measure the mobility and interactions of single proteins in living cells.^[3–7]^ In an SPT experiment, the movement of fluorophore-labeled single proteins is measured, and mobility trajectories are generated, providing spatiotemporally-resolved information on diffusion properties.^[8]^ A variety of methods for fluorophore labeling was established, including fluorescent proteins,^[9]^ fluorophore-labeled ligands^[10,11][12]^ or nanobodies,^[13,14]^ as well as self-labeling protein tags such as HaloTag^[15–17]^ or SNAP-tag^[18,19]^. The integration of photoactivatable fluorescent proteins allowed SPT at high protein densities and allowed for high-density mapping of protein mobility.^[20]^

As with any fluorescence method, SPT is challenged by photobleaching, which affects both the length of trajectories as well as the observation time for a single cell.^[3]^ The trajectory length is mainly limited due to the photon budget that is available from a single fluorophore. More photostable fluorescent probes, such as quantum dots, provide longer trajectories, yet might pose a challenge because of their size and more challenging conjugation chemistry.^[21]^ Recently, the implementation of DNA as a protein label, conjugated to protein-targeting nanobodies, was shown to enable enhanced observation times.^[22,23]^ The limit in observation time of a single cell is a consequence of using high-affinity or covalent protein labels and ultimately photobleaching. This can be bypassed by using weak-affinity fluorophore labels, which continuously bind to and unbind from a protein target, from a reservoir.^[24]^ For example, retro-engineered HaloTags designed for reversible substrate exchange enable longer observation times.^[25]^

Tracking single-molecule diffusion for extended observation times and in single cells is needed to monitor membrane receptor activation. SPT is an ideal method, as it allows to observe these processes in the native environment of a live cell, and since diffusion properties can serve as a proxy readout for membrane protein activation. However, ligand-induced activation of membrane receptors occurs at the time scale of several minutes.^[26,27]^ Because of this, SPT experiments monitoring membrane receptor activation were so far conducted by sequential single-molecule imaging of many single cells.^[13,28]^

Here, we report that the self-labeling protein tag HaloTag7 in combination with recently developed exchangeable HaloTag Ligands (xHTLs)^[29]^ enables single-particle tracking in single living cells for extended observation times. First, we compared the fluorescence signal over time of xHTLs to covalent HaloTag7 ligands and observed an almost continuous fluorescence signal for xHTLs in live-cell SPT experiments. Next, we demonstrated robust single-molecule diffusion analysis of xHTL-labeled membrane proteins in live cells. We then demonstrate long-time SPT experiments of up to 30 minutes in single living cells expressing a HaloTag7-fusion of the epidermal growth factor receptor (EGFR), before and after activating with its cognate ligand EGF. We found that the temporal change of diffusion properties mirrors the previously reported activation times of EGFR.^[26]^ Furthermore, long-time SPT allowed to generate a spatial map of diffusion in single cells, and revealed locally confined areas of EGFR activation in the plasma membrane.

## Results

The self-labeling protein tag HaloTag7 (HT7) in combination with exchangeable HaloTag Ligands (xHTLs) that repetitively bind to and unbind from a protein target was shown to provide a constant fluorescence signal in various fluorescence microscopy applications and excellent compatibility with live-cell imaging.^[29–31]^ We, therefore, reasoned that this labeling strategy should enable long observation times in SPT experiments in single living cells.

To evaluate this concept, we fused HT7 to the intracellular domain of CD86, a monomeric membrane protein,^[32]^ and generated U-2 OS cell lines stably expressing CD86-HT7 (**Figure 1A**). We measured the mobility of single CD86-HT7 proteins in living cells and generated single-molecule trajectories. We then compared the density of single-molecule trajectories in living cells for a non-covalent xHTL (SiR-S5) to a covalent HTL (SiR-HTL) and found a strong decrease for the covalent SiR-HTL already after 10 s (**Figure 1BC**). The dead-mutant dHT7 (D106A), an orthogonal protein tag to HT7,^[29]^ was labeled with xHTL SiR-Hy5 and showed very similar preservation of the fluorescence signal over time (**Figure 1C**). Nonspecific binding of all tested HT ligands to the plasma membrane was low in control experiments in U-2 OS wild-type cells that do not express a HaloTag7 fusion protein (**Figure S1**).

**Figure 1.**
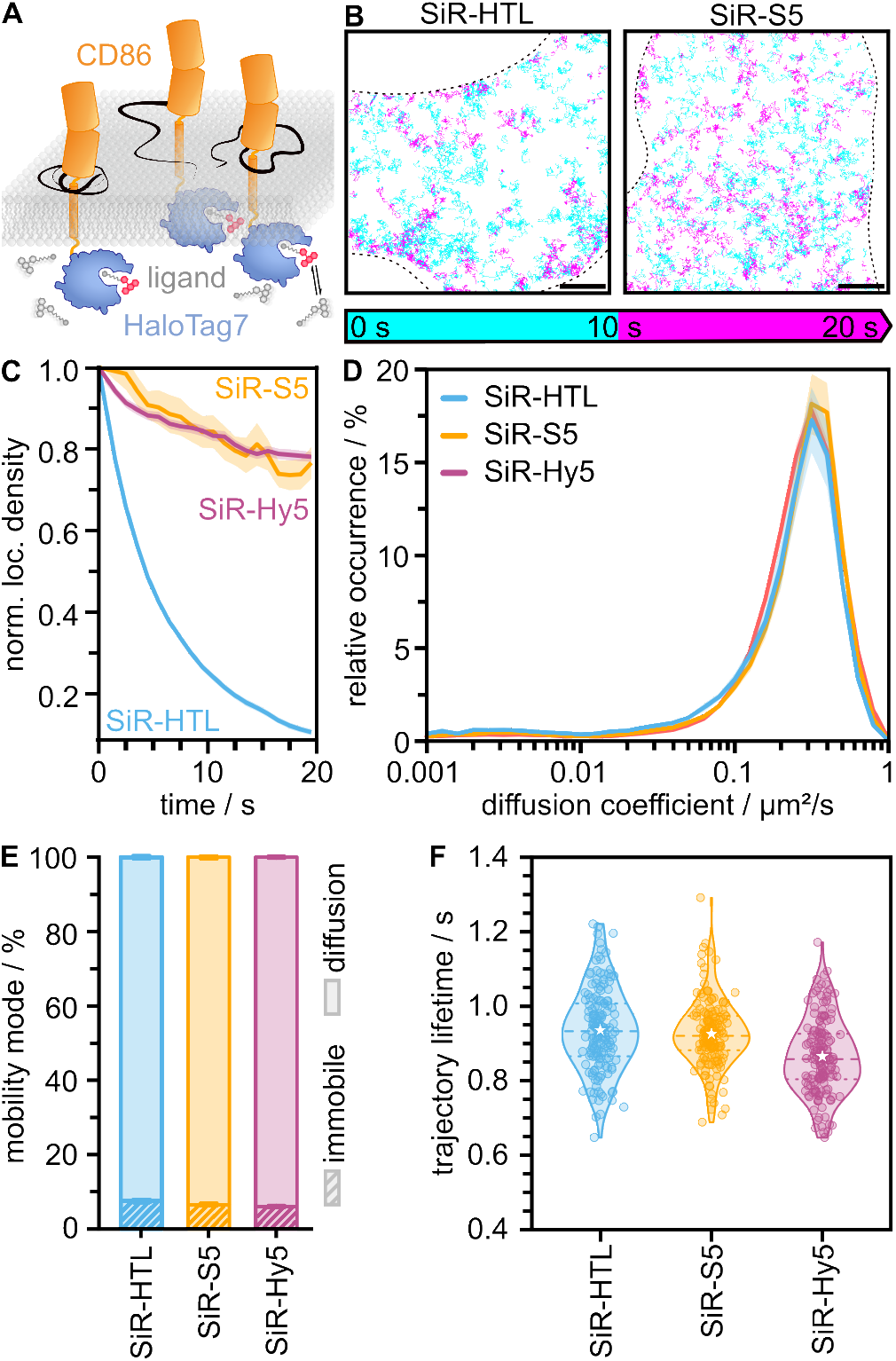
Single-particle tracking in live cells using exchangeable HaloTag Ligands (xHTLs). (A) Principle of xHTL-SPT shown for the transmembrane protein CD86-HT7. Fluorogenic xHTLs reversibly bind to and unbind from the HT7 tag fused to the intracellular region of CD86 and enable single-molecule tracking. (B) Single-molecule trajectories of exemplary cells acquired using the covalent SiR-HTL (1 nM) or xHTL SiR-S5 (1 nM) as probes on CD86-HT7. Trajectories are color-coded based on their first appearance in the first (cyan) or later half (magenta) of the 20 s acquisition time. Scale bar 5 µm. (C) Mean number of localizations per area binned into 1 s intervals, normalized to the respective data in the first frame, and plotted over time for HT7/SiR-HTL (blue), HT7/SiR-S5 (yellow), and dHT7/SiR-Hy5 (magenta). (D) Relative occurrence of the mean diffusion coefficient per cell for SiR-tagged exchangeable HaloTag Ligands and the covalent SiR-HTL. (E) Percentage of mobility modes per cell for SiR-HTL, SiR-S5, and SiR-Hy5 (1 nM each). Single-molecule trajectories were assigned to the classes immobile or diffusion. (F) Lifetime of single-molecule trajectories of the covalent SiR-HTL and exchangeable SiR-S5 binding to HaloTag7 and SiR-Hy5 binding to dHaloTag7. Dashed lines represent the median, stars the mean, and dotted lines the interquartile range. All errors represent the standard error of the mean (SEM). 160 cells were acquired for each condition in (C)-(F).

To evaluate the xHTL-HT7/-dHT7 labeling concept for diffusion analysis, we compared the diffusion coefficient and diffusion mode of CD86 fusions to covalently labeled CD86-HT7. Independent of the HaloTag-ligand combination used, the distribution of diffusion coefficients as well as the mobile fraction were similar, yielding mean diffusion coefficients of approx. 0.3 µm^2^/s and about 7% immobile particles (**Figure 1DE, Table S1, Table S2**). We next determined the trajectory lifetime SiR-labeled ligands and found reasonably similar values of 0.94 ± 0.03 s (SiR-HTL), 0.93 ± 0.04 s (SiR-S5) and 0.87 ± 0.02 s (SiR-Hy5) (**Figure 1F, Table S3**).

We next evaluated the influence of the fluorophore on the diffusion analysis. For this purpose, we used xHTLs conjugated to the fluorophore JF_585_, measured the diffusion properties of CD86-HT7/-dHT7, and found very similar diffusion coefficients of approximately 0.3 µm^2^/s (**Figure S2A, Table S1**). We found some small variations in the fraction of immobile particles for JF_585_-S5 and JF_585_-Hy4, as compared to SiR-xHTLs (**Figure S2B, Table S2**). The trajectory lifetime of JF_585_-xHTLs was found to be shorter (0.6 s) than those of SiR-labeled xHTLs (**Figure S2C, Table S3**). The fluorescence signal over time remained almost constant, indicating a continuous exchange of xHTL at the HaloTag (**Figure S2D, Table S3**). These results indicate that diffusion coefficients and diffusion modes are independent of the fluorophores tested and the type of HT ligand. We also fused HT7 or dHT7 to the intracellular domain of CTLA-4, a dimeric membrane protein,^[33]^ and performed SPT experiments in live U-2 OS cells stably expressing the fusion protein. Again, we observed only a minor decrease of the fluorescence signal over time, and diffusion properties of CTLA-4 fusions were found to be similar for a variety of xHTL-fluorophore/HT7 combinations (**Figure S3**). Taken together, the xHTL-HT7/-dHT7 system in combination with SPT experiments provides robust information on protein diffusion, while bringing in the additional benefit of minimal signal loss over time.

We observed that xHTL labeling generates a homogeneous distribution of single-molecule trajectories in cells, whereas HTL labeling leads to more trajectories at the cell border (**Figure 1B**). To quantify this observation, we measured the fluorescence signal in living cells at the cell border and compared it to the remaining cell surface area (**Figure S4**). For the exchangeable probe SiR-S5, the signal was constant throughout the cell surface area, whereas for the covalent ligand SiR-HTL, the fluorescence signal was increased at the cell border, indicating a higher protein concentration. We attribute this increase in fluorescence intensity at the cell border to photobleaching of covalent HTLs in the field of imaging of the basal membrane, and replenishment of HTL-HT7 through diffusion from the apical membrane across the cell borders (**Figure S4**).

Motivated by the preservation of fluorescence signal and trajectory density, we attempted to establish long observation times in single living cells by using xHTL/HT7 for repetitive and continuous labeling of target proteins. We conducted long-term SPT experiments of HT7/dHT7-tagged CD86 in live cells for up to 20 min using the xHTLs SiR-S5 and SiR-Hy5. We found that CD86-HT7 labeled with the exchangeable SiR-S5 yielded an almost constant trajectory density in living cells, whereas, with the covalent SiR-HTL, the density dropped off quickly (**Figure 2AB**). This finding is corroborated by the single-molecule signal detected over time, which quickly decreases for the covalent SiR-HTL, and remains at a higher level (SiR-Hy5) or even almost constant (SiR-S5) for the exchangeable ligands (**Figure 2, Figure S3E**). These findings qualify exchangeable HaloTag7 ligands for long-term SPT experiments in living cells and spatial mapping of the diffusion landscape.

**Figure 2.**
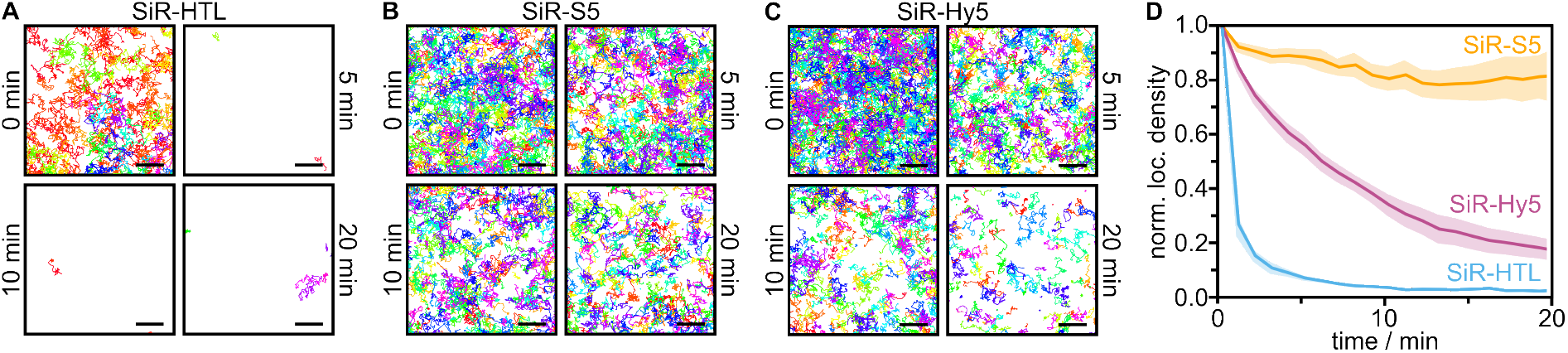
Long-time single-particle tracking of CD86-(d)HT7 in living U-2 OS cells. (A, B, C) Single-molecule trajectories of CD86 in exemplary cells obtained from (A) HT7/SiR-HTL (covalent), (B) HT7/SiR-S5, or (C) dHT7/SiR-Hy5 (exchangeable). Probe concentration was 1 nM. For each snapshot, the trajectories occurring within 1 min are shown (scale bars 2 µm). (D) Mean number of localizations per area binned into 1 min intervals, normalized to the respective data in the first frame, and plotted over time for CD86-HT7 or CD86-dHT7 imaged with SiR-HTL (blue, N = 5), SiR-S5 (yellow, N = 5), and SiR-Hy5 (magenta, N = 6) (1 nM ligand concentration). Error bands represent the SEM.

Receptor mobility can serve as a readout to monitor functional states. In cells stimulated with a receptor-targeting ligand, a slower diffusion, or immobilization, can be correlated with activation.^[34]^ Since the activation occurs at a timescale of several minutes,^[26]^ the accessible observation time of single-cell SPT experiments was so far a limitation to monitor receptor activation in individual cells. This has been bypassed by sequential imaging of several cells and temporal tiling of SPT data.^[13,28]^ We reasoned that xHTL/HT7 labeling should provide a direct solution to monitor receptor activation.

We measured the diffusion properties of the transiently expressed epidermal growth factor receptor (EGFR) before and after stimulation with its native ligand EGF in single living U-2 OS cells. For this purpose, we fused HT7 to the C-terminal (intracellular) site of EGFR, labeled it with SiR-S5, and conducted SPT experiments in living U-2 OS cells. First, we calculated the diffusion coefficient and mode of EGFR from 20 s measurements in resting and EGF-stimulated cells. We observed a reduction in the diffusion coefficient (from 0.245 ± 0.004 µm^2^/s to 0.158 ± 0.003 µm^2^/s) and an increase in immobile EGFR receptors upon EGF treatment (from 6.9 ± 0.2% to 10.1 ± 0.4%) (**Figure 3ABC, Figure S5, Table S2**). These findings are in line with previously published work^[34,35]^ and the model of EGFR activation.^[36]^

**Figure 3.**
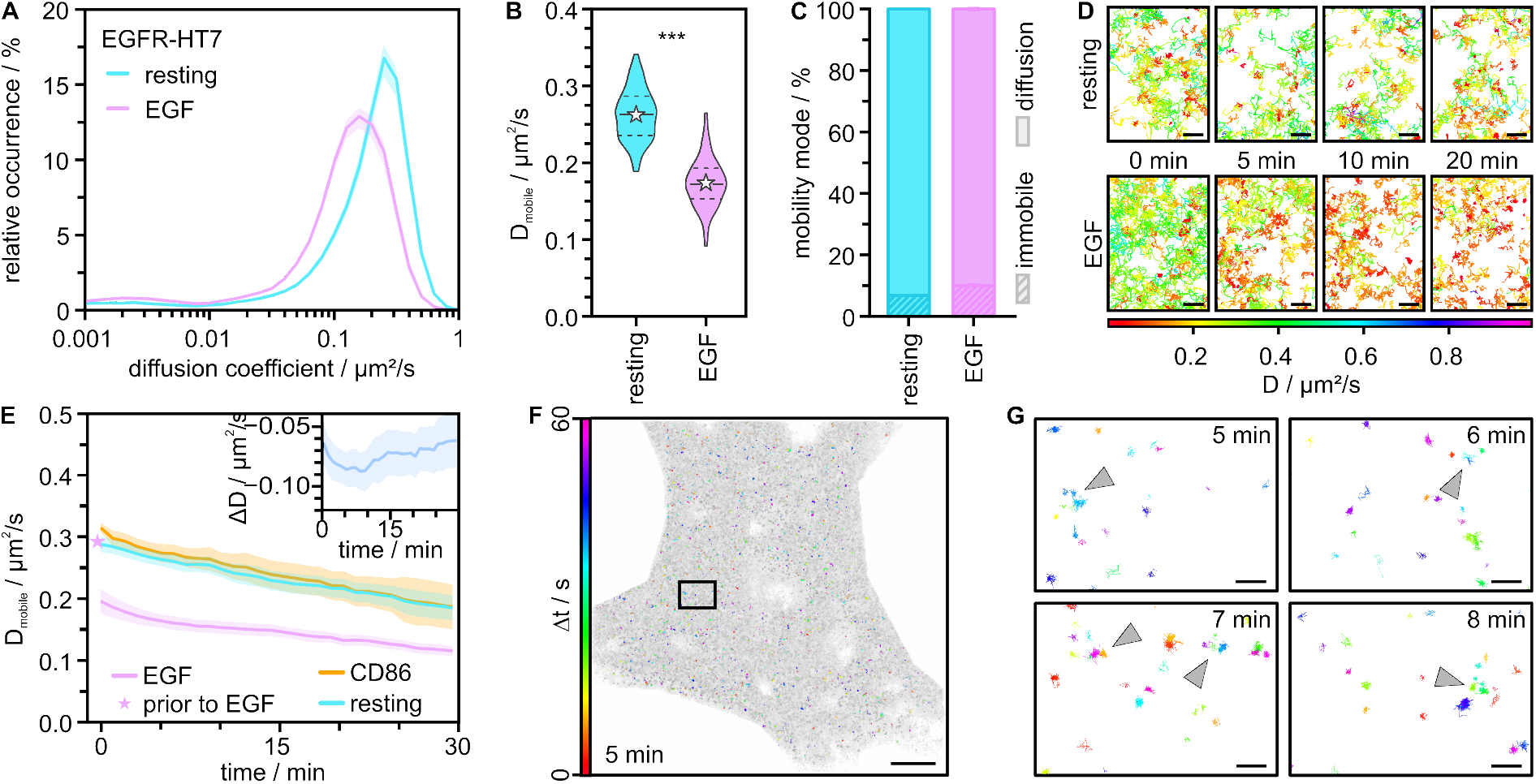
Long-time SPT of transiently expressed EGFR-HT7 in living U-2 OS cells measured with 1 nM SiR-S5. (A) Relative occurrence of the mean diffusion coefficient per cell for resting (cyan) and EGF-treated (20 nM) cells (magenta). (B) Distribution of diffusion coefficients of mobile EGFR-HT7 extracted from resting and EGF-treated cells. Dashed lines represent the median, stars the mean, and dotted lines the interquartile range. P < 0.001 (***) highly significant difference between means. (C) Percentage of mobility modes per cell for EGFR in unstimulated and EGF-stimulated cells. Single-molecule trajectories were assigned to the classes immobile or diffusion. Data in A-C were taken from 20 s measurements (N = 120). (D) Single-molecule trajectories detected at different time points of a long-time measurement for resting and EGF-stimulated cells color-coded by the assigned diffusion coefficient. (E) Mean diffusion coefficient of mobile trajectories binned into 1 min intervals and plotted over time for CD86-HT7 labeled with SiR-S5 (orange, N = 4), EGFR-HT7 imaged with SiR-S5 in resting (cyan) and EGF-treated cells (magenta) (N = 4). Stars represent the diffusion coefficient in cells before EGF stimulation. The inlay shows the difference plot for the diffusion coefficient of EGFR-HT7 in EGF-treated cells. All errors represent the standard error. (F) All trajectories detected over the 30 min acquisition time (gray) overlaid by the immobile trajectories from the period of 300-360 s (color-coded by their time of appearance within one minute, rainbow) (scale bar 5 µm). (G) Zoom-in of the box in (F) at different time points (5, 6, 7, and 8 min) of the measurement. Immobile trajectories are clustering in local areas (indicated by arrowheads). Trajectories are color-coded by the time of appearance within one minute (scale bar 500 nm).

We next performed SPT experiments of EGFR-HT7 in living cells over an extended observation time of up to 30 min and analyzed its diffusion properties. In untreated cells, we observed some continuous decrease in diffusion coefficients throughout the acquisition time of 30 min. In EGF-stimulated cells, the fraction of slower diffusing EGFR-HT7 increased strongly over time (**Figure 3D**). We quantified the diffusion coefficient of mobile EGFR-HT7 over time in untreated cells and EGF-treated cells and compared it to CD86-HT7. We found that EGFR-HT7 and CD86-HT7 show a continuous decrease in diffusion coefficient over 30 min, which might be a result of minor phototoxic effects during long-time fluorescence imaging in living cells (**Figure 3E**). However, the decrease in the diffusion coefficient was much stronger for EGFR-HT7 in EGF-treated cells, particularly in the first 10 min. To extract the activation kinetics of EGFR, we used the diffusion profile of untreated EGFR-HT7 as a baseline and subtracted it from the diffusion profile of EGFR-HT7 in EGF-treated cells. This delivered a difference plot for the diffusion coefficient of untreated and EGF-stimulated EGFR-HT7, which showed a dip at around 5 to 10 mins, followed by a recovery (**Figure 3E, inset**).

Measuring the diffusion of EGFR-HT7 in single EGF-treated cells for extended observation times also allowed us to spatially map membrane sites where receptor activation occurs (**Figure 3FG, Figure S6**). We found a non-uniform distribution of immobile EGFR-HT7 at different time points, and the appearance of local ‘hot spots’ that accumulated several receptors at short distances for a short time (**Figure 3G**, arrowheads).

The two orthogonal HaloTag variants open the possibility for dual-color labeling of two different proteins. In combination with exchangeable ligands, long-term dual-color SPT is possible. To demonstrate this, EGFR-HT7 and CD86-dHT7 were co-transfected into U-2 OS cells (**Figure 4A**) and simultaneously tracked in live cells using the two orthogonal exchangeable ligands SiR-S5 and JF585-Hy4 (**Figure 4B**).

**Figure 4.**
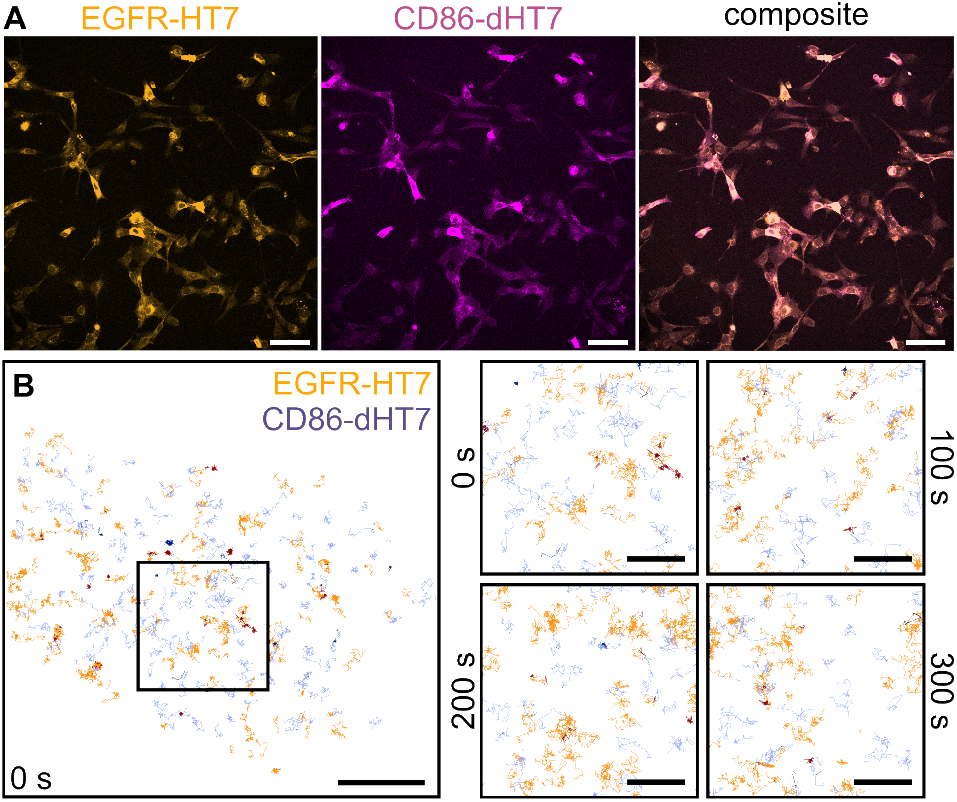
Dual-color SPT of two orthogonal HaloTags. (A) Confocal microscopy images of U-2 OS cells transiently expressing EGFR-HT7 labeled with 100 nM SiR-S5 (orange) and CD86-dHT7 labeled with 100 nM JF_585_-Hy4 (magenta) together with the composite of both channels (scale bars 100 µm). Nearly all transfected cells express both fusion proteins. (B) Single-molecule trajectories reconstructed from 20 s of an SPT experiment for EGFR-HT7 (orange) and CD86-dHT7 (purple) labeled with 1 nM SiR-S5 and JF_585_-Hy4, respectively (scale bar 5 µm). Mobile trajectories are represented in light colors, immobile trajectories in dark colors. The box marks the region shown as zoom-ins at different time points on the right (scale bars 2 µm).

## Discussion

SPT experiments with covalent or high-affinity protein labels enable observation times of seconds to a few minutes.^[7,35,37–39]^ Here, we introduce the self-labeling protein tag HaloTag7 in combination with weak-affinity and exchangeable fluorophore ligands (xHTLs) for SPT experiments with extended observation times in single living cells. In SPT experiments with HT7 fused to membrane proteins, we compared the performance of HT7 labeled with xHTLs to covalent HTLs. We found that xHTLs show similar trajectory lengths and diffusion properties as HTLs for the fluorophore SiR (**Figure 1BC)** while enabling largely extended observation times in single cells (**Figure 2, Figure 3**). We employed fluorogenic xHTLs, reducing the background signal and enhancing the signal-to-background ratio.^[29]^

A relevant parameter for SPT is the trajectory length, which determines the accuracy of the diffusion analysis through the mean-squared displacement (MSD).^[40,41]^ The trajectory lifetime of xHTLs determined in this work is similar to that of fluorophore-labeled nanobodies or ligands.^[13,14,42]^ The trajectory length of xHTLs depends on i) fluorophore photobleaching, ii) photodamage to the HaloTag7 protein which might make it unable to efficient ligand binding, and iii) the binding time to the HT7/dHT7 protein. SiR-S5, SiR-Hy5 and SiR-HTL show similar trajectory lengths of 0.87 - 0.94 s (**Figure 1F, Table S3**), while previously reported binding times of xHTLs are around 2 s (SiR-S5) and 1 s (SiR-Hy5). This indicates that the trajectory lifetime of SiR-S5 is more determined by fluorophore photobleaching or photodamage of the protein tag, while the trajectory lifetime of SiR-Hy4 is also influenced by the shorter binding time to dHT7. The trajectory length observed for JF_585_-xHTLs was shorter than for SiR-xHTLs, indicating that the major contribution here is fluorophore photobleaching. The xHTL/HT7/dHT7 system complements previous work on retro-engineered HaloTag-HTL that were designed for longer binding times, yet with a similar accuracy in diffusion analysis.^[25]^ Yet, the xHTL/HT7 labeling pair profits from the availability of engineered cell-lines with HT7-fusion proteins. DNA-labeled protein ligands enable even longer trajectory lifetimes but are limited to extracellular labeling that might impair ligand-binding.^[23]^

The transient and repetitive binding of xHTL to HT7-tagged proteins allows continuous SPT imaging in living cells for extended observation times. This enables monitoring the diffusion of membrane receptors in single cells over time periods that match receptor activation kinetics, i.e. several tens of minutes, and which could previously only be accessed by measuring multiple cells sequentially^[13,28]^ or limited to extracellular target sites.^[23,43]^ To demonstrate the potential of the xHTL/HT7 labeling pair, we conducted SPT experiments with EGFR-HT7 in live cells. We chose EGFR as it is a membrane receptor that has been extensively studied with various SPT methods.^[44]^ The diffusion coefficient we measured in resting and EGF-treated cells (0.245 ± 0.004 µm^2^/s and 0.158 ± 0.003 µm^2^/s) are within the range of reported values (0.02-0.5 µm^2^/s and 0.005-0.29 µm^2^/s).^[34,35,39,45–51]^ In addition, our approach enabled us to follow the EGF-induced activation kinetics of EGFR directly in single living cells, by monitoring the diffusion coefficient and diffusion mode (**Figure 3E**). With that, we extracted for the first time the activation kinetics of EGFR following EGF-treatment directly in a single living cell, peaking at a time of 5-10 minutes after ligand stimulation and matching cell biology data (**Figure 3E inset**).^[26,27]^ Extended observation times also enabled the observation of local accumulations of immobile EGFR in spatially confined ‘hot-spots’ of sub-micrometer size (**Figure 3G**). Such ‘hot spots’ indicate the presence of membrane regions where signaling hubs build preferentially and endocytosis through clathrin-coated pits might occur.^[34,52]^ Similarly sized ‘hot spots’ were found in diffraction-limited microscopy in live cells.^[48]^ A non-uniform distribution of receptor activation was also reported for the EGFR/HER2 heterodimer.^[53]^ In summary, the presented method qualifies SPT to monitor membrane receptor activation in single living cells at extended observation times matching the kinetics of ligand-receptor activation.^[26,27]^ We further showed dual-color SPT in live cells with two orthogonal HaloTags in combination with exchangeable ligands. The xHTL-SPT approach allows simultaneous tracking of different proteins over extended periods (**Figure 4B**).

A challenge of any long-time fluorescence microscopy experiment in living cells is the potential accumulation of phototoxic molecules, which might affect cell viability. This might in part explain the observed decrease in diffusion coefficient in long-term experiments found for all membrane receptors investigated in this work (**Figure 3E**).^[54]^ Albeit that a correction with a baseline was possible and activation kinetics of EGFR could be extracted, it would be interesting to explore strategies to mitigate these effects. One possible solution might be reduced laser intensities in combination with content restoration through e.g. denoising.^[55,56]^ Another solution might be fluorophores that are designed or manipulated such that they less often enter phototoxicity-inducing photophysical states.^[57–61]^

Another advantage of using exchangeable fluorophore labels in live-cell microscopy of membrane surface proteins is a spatially homogeneous fluorescence signal. This is in contrast to covalent protein labels, which photobleach in the illumination area and are replenished from the border regions of a cell, leading to apparently higher protein densities at cell edges (**Figure 1B, Figure S4**) that are often seen in SPT experiments.^[10,28,62,63]^ The observation of signal accumulation at cell borders appears to be generally valid for live-cell microscopy of membrane proteins labeled with covalent labels (**Figure S4**).

## Conclusion

The xHTL/HT7 labeling pair enables SPT experiments in live cells with extended observation times of up to 30 minutes. Due to its versatility, use of standard protein tags, range of ligands and fluorophores as well as ease of use, this method has the potential to become a valuable tool for studying long-term protein dynamics in live cells with single-molecule resolution. The approach is transferable to other proteins and opens the window to study receptor signaling kinetics in live cells.

## Materials and Methods

### Fluorescent probes

All fluorescent xHTLs (SiR-S5, SiR-Hy5, JF_585_-S5, and JF_585_-Hy4) were prepared as described in Kompa *et al*.^[64]^ SiR-HTL was kindly provided by B. Réssy and synthesized according to the literature procedure.^[65]^

### Plasmids

HaloTag7 (HT7) or the dead-mutant of the HaloTag7 (dHT7) was genetically fused to the C-terminus of CD86 (Addgene plasmid #98284),^[66]^ CTLA-4 (Addgene plasmid #98285)^[66]^ and EGFR (Addgene plasmid #32751)^[67]^ by replacing the fluorescent protein (mEos2 or EGFP) by molecular cloning (Gibson assembly^[68]^). For expression in mammalian cells, the fusion construct was subcloned into a pcDNA5/FRT/TO vector (Invitrogen, Thermo Fisher Scientific, Waltham, MA, USA). All plasmid sequences were verified by Sanger sequencing (Microsynth AG, Balgach, Switzerland).

### Generation of stable cell lines

Stable cell lines were generated using the Flp-IN T-REx^TM^ system (Invitrogen). In brief, U-2 OS Flp-In T-REx cells^**[69]**^ were co-transfected with a 1:10 ratio of pcDNA5-FRT-TO-GOI and pOG44 (Invitrogen) plasmids using the Lipofectamine3000 reagents (Invitrogen) according to the manufacturer’s protocol. On the next day, cells were selected with 100 μg/mL hygromycin B (Gibco, Thermo Fisher Scientific, Waltham, MA, USA) for 2 days. After recovery, the cells were treated with 100 μg/mL doxycycline (Sigma-Aldrich, St. Louis, MO, USA) for 24 h to induce protein production previous to fluorescence-assisted cell-sorting (FACS) on a Melody^TM^ Cell sorter (BD Biosciences, Franklin Lakes, NJ, USA) for transgene expression.

### Cell culture

U-2 OS wild-type cells (CLS Cell Lines Service GmbH, Eppelheim, Germany) and all stable cell lines were cultivated in growth medium (Dulbecco’s Modified Eagle Medium: Nutrient Mixture F12 without phenol red (Gibco) supplemented with 1% GlutaMAX (Gibco), 10% FBS (Sartorius, Göttingen, Germany), 100 U/mL penicillin (Gibco), and 100 µg/mL streptomycin (Gibco)) at 37 °C and 5% CO_2_ in an automatic CO_2_ incubator (Model C150, Binder GmbH, Tuttlingen, Germany).

For live-cell microscopy, cells were seeded to a density of 4×10^4^ cells per sample onto PLL-PEG-RGD-functionalized coverslips in 6-well plates as described elsewhere.^[11]^ After 2 d incubation, protein expression was induced with 100 ng/mL doxycycline (Sigma-Aldrich). Cells were incubated for a further day before performing live-cell microscopy experiments.

For live-cell imaging of EGFR-HT7, the respective plasmid was transiently transfected into U-2 OS wild-type cells. 35×10^4^ cells were seeded in 6-well plates, transfected 2 d later with 1 µg/well plasmid DNA using Lipofectamin LTX (Invitrogen) according to the manufacturer’s protocol, and incubated at 37 °C and 5% CO_2_. After 6 h, cells were transferred onto PLL-PEG-RGD-coated coverslips via trypsinization and split 1:3. The transferred cells were incubated for 1 d on the coated coverslips in growth medium at 37 °C and 5% CO_2_ before imaging.

For confocal and two-color live-cell imaging of EGFR-HT7 and CD86-dHT7, the respective plasmids were transiently co-transfected into U-2 OS wild-type cells. 30×10^4^ cells were seeded in 6-well plates, transfected after 2 d with 1 µg/well per plasmid using jetOPTIMUS transfection reagent (Polyplus, Illkirch, France) according to the manufacturer’s protocol, and incubated at 37 °C and 5% CO_2_. After 7 h, cells were transferred onto PLL-PEG-RGD-coated coverslips via trypsinization and split 1:2. The transferred cells were incubated for 1 d on the coated coverslips in growth medium at 37 °C and 5% CO_2_ before imaging.

### Sample preparation

For live-cell imaging using covalently binding HaloTag Ligands (HTLs), cells were stained with 1 nM SiR-HTL in growth medium for 30 min at 37 °C and 5% CO_2_ prior to experiments. Then, coverslips were installed into custom-built holders and rinsed three times with FluoroBrite DMEM (Gibco), also used as imaging medium. For live-cell imaging with exchangeable HaloTag Ligands (xHTLs), coverslips were directly mounted into the holders, covered with FluoroBrite DMEM as imaging medium, and supplemented with 1 nM ligand (SiR-S5, SiR-Hy5, JF_585_-S5 or JF_585_-Hy4, introduced in Kompa *et al*.^[29]^). Samples were mounted into a stagetop incubator (Okolab, Otaviano, Italy) at 25 °C for 10 min before measurements.

For EGF-stimulated samples, the ligand (PeproTech, Thermo Fisher Scientific, Waltham, MA, USA) was added to the sample directly on the microscope to a final concentration of 20 nM (see below).

### Live-cell microscopy

Single-color live-cell microscopy experiments were conducted at 25 °C on a commercial widefield microscope (N-STORM; Nikon, Düsseldorf, Germany). The system was controlled via NIS Elements (v4.30.02, Nikon), and LCControl (Agilent, Santa Clara, California, USA), and equipped with an oil-immersion objective (100×Apo TIRF oil, NA 1.49), a stage top incubator (Okolab, Pozzuoli, Italy) and an EMCCD camera (Andor iXon, DU-897U-CS0-BV, Andor, Belfast, UK), operated in total internal reflection fluorescence (TIRF) mode. Samples were illuminated by a 561 nm at 0.5 kW/cm^2^ or 647 nm laser at 0.4 kW/cm^2^. For long-term measurements of EGFR, a reduced 647 nm laser power of 0.06 kW/cm^2^ was used. The following camera parameters were set: EM gain 200, pre-amplifier gain 3, read-out rate 17 MHz, and active frame transfer. Image stacks of 256×256 px with a pixel size of 157 nm were recorded with µManager (v1.4.22)^[70]^ at an integration time of 20 ms. Either 1,000 (standard measurements), 60,000 (long-term measurements of CD86 and CTLA-4), or 90,000 frames (long-term measurements of EGFR) were recorded per cell, while each sample was imaged for a maximum of 30 min. For long-term measurements of EGF-stimulated samples, a cell was recorded for 1,000 frames in resting condition, then 20 nM EGF were added, and the same cell was imaged for 90,000 frames.

Two-color live-cell microscopy experiments were performed on a home-built Olympus IX-71 inverted TIRF microscope (Olympus Deutschland GmbH, Hamburg, Germany) equipped with a nosepiece stage (IX2-NPS, Olympus Deutschland GmbH) to provide z-plane adjustment and minimization of drift during measurements. Two lasers (561 nm, 200 mW Sapphire and 637 nm, 140 mW OBIS, both Coherent Inc., Santa Clara, CA, USA), colinearly superimposed using a dichroic mirror (H 568 LPXR superflat, AHF Analysentechnik AG, Tübingen, Germany), served as excitation sources. The sample was illuminated with both lasers after passing an acousto-optical tunable filter (AOTF; AOTFnC-400.650-TN, AA Opto-Electronic, Orsay, France). The two lasers were coupled by a fiber collimator (PAF-X-7-A, Thorlabs, Dachau, Germany) into a single-mode optical fiber (P5-460AR-2, Thorlabs) and subsequently re-collimated to a diameter of 2 mm (60FC-0-RGBV11-47, Schäfter & Kirchhoff, Hamburg, Germany). The collinear beams were directed to a 2-axis galvo scanner mirror system (GVS012/M, Thorlabs) where electronic steering, controlled by an in-house Python script, allowed switching between illumination modes. The excitation light passed two telescope lenses (AC255-050-A-ML and AC508-100-A-ML, Thorlabs) focusing them onto the back focal plane of the objective (UPlanXApo, 100x, NA 1.45, Olympus Deutschland GmbH). In a filter cube, which directs the beam into the objective, two clean-up and rejection bandpass filters together with a dichroic mirror were installed (Dual Line Clean-up ZET561/640x, Dual Line rejection band ZET 561/640, Dual Line beam splitter zt561/640rpc, AHF Analysentechnik AG). Fluorescence light was collected through the same objective and passed the dichroic mirror toward the detection path. An Optosplit II (Cairn Research Ltd, UK) split the emission light around 643 nm into two channels with a beam splitter and two bandpass filters (H643 LPXR, 605/52 BrightLine HC, 679/41 BrightLine HC, AHF Analysentechnik AG). The spatially separated SiR and JF_585_ channels were simultaneously detected on an EMCCD camera (iXon Ultra X-10971, Andor Technology Ltd, Belfast, UK). JF_585_ and SiR were excited with 0.1 kW/cm^2^ (637 nm) or 0.05 kW/cm^2^ (561 nm), respectively, in circular TIRF mode. The following camera parameters were set: EM gain 200, pre-amplifier gain 3, read-out rate 17 MHz, and active frame transfer. Image stacks of 256×256 px with a pixel size of 159 nm were recorded with µManager (v2.0.0)^[70]^ at an integration time of 20 ms.

For each sample, 10 background measurements with 1,000 frames were taken in areas without cells.

### Confocal microscopy

Samples were prepared as described above for live-cell microscopy, but the imaging medium was supplemented with 100 nM SiR-S5 and JF_585_-Hy4. Confocal microscopy experiments were conducted at room temperature on a commercial SP8 confocal laser scanning microscope (Leica Microsystems, Wetzlar, Germany) equipped with an HC PL APO CS2 20x/0.75 immersion objective (Leica Microsystems). Images were acquired with the Leica Application Suite X Software (v3.5.7.23225, Leica Microsystems). 16-bit images with a size of 1024 × 1024 px at 757.58 nm pixel size were acquired with a scan speed of 400 Hz and a line average of 2. SiR was excited with a 633 nm HeNe laser at 0.2% intensity and its emission was detected with a HyD detector (gain 160) in a 638-784 nm detection window. JF_585_ was excited with a 561 nm HeNe laser at 0.5% intensity and its emission was detected with a HyD detector (gain 160) in a 573-628 nm detection window.

### Data analysis

SPT data were analyzed using a pipeline for single-particle tracking analysis as described in detail elsewhere.^[11,13,42]^ In brief, raw data were localized using ThunderSTORM (dev-2016-09-10-b1),^[71]^ a plugin for Fiji,^[72]^ localizations were connected to trajectories in swift (v0.4.3),^[73]^ and parameter estimation for swift as well as the diffusion analysis was conducted in SPTAnalyser (v.1.2.0).^[42]^ Parameters were determined according to the SPTAnalyser software manual (v1.2.0) and chosen as published previously,^[13]^ if not stated otherwise in the following. For SiR-labeled ligands, the parameter *diffraction_limit* was set to 18 nm, *exp_displacement* to 125 nm, *p_bleach* to 0.025, and *D*_*min*_ to 0.0041 µm^2^/s. For JF_585_-labeled ligands, *diffraction_limit* = 15 nm, *exp_displacement* = 135 nm, *p_bleach* = 0.08 nm, and *D*_*min*_ *=* 0.0057 µm^2^/s was applied.

SPTAnalyser including scripts for batch processing is available on github (https://github.com/HeilemannLab/SPTAnalyser) together with detailed documentation.

For long-term measurements, the localizations per frame were extracted from the localization lists, normalized to the area of the cell, binned into 1 min (3,000 frames) intervals, and normalized to the first frame. The diffusion coefficients per frame were directly extracted from the output files from swift, and filtered for mobility type and trajectory length. Only diffusion coefficients of trajectories classified as mobile and longer than 20 frames were kept. For every trajectory, only a single diffusion coefficient was counted in the frame in which the respective trajectory appeared. To visualize the activation profile of EGFR, the diffusion coefficients per frame of EGF-treated cells were binned into 1 min intervals and subtracted by the diffusion coefficients per frame of resting cells, to yield the difference in diffusion coefficients ΔD.

All errors represent the standard error of the mean (SEM). OriginPro 2024 (v10.1.0.170, OriginLab Corporation, Northampton, MA, USA) was used for statistical analysis. The Shapiro-Wilk test (α = 0.05) was applied to test populations for normality. The Mann-Whitney U test was applied for statistical testing as some data sets rejected normality. p-values < 0.001 are marked as highly significantly different (***).

## Acknowledgments

We are thankful to Dr. Maja Klevanski (Heidelberg University) for helpful discussions. CC, MSD, SJ, PF, MH were supported by the Deutsche Forschungsgemeinschaft (DFG, German Research Foundation) through SFB 1507 (project-id: 450648163) and DFG INST 161/778-1 FUGG. MH and SJ acknowledge funding by the International Max-Planck Research School on Cellular Biophysics (IMPRS-CBP). JK and KJ were supported by the Max Planck Society, the Ecole Polytechnique Fédérale de Lausanne (EPFL), and the Deutsche Forschungsgemeinschaft (DFG, German Research Foundation), TRR 186.

## Author contributions

MH designed the research. CC, MSD, JK, SJ, and PF performed experiments. CC and MSD contributed to data analysis. CC, MSD, JK, and MH wrote the manuscript with contributions from all authors.

## References

[1] P. L. Yeagle, in The Membranes of Cells (Ed.: P.L. Yeagle), Academic Press, 2016.

[2] C.-H. Heldin, B. Lu, R. Evans, J. S. Gutkind, Cold Spring Harb. Perspect. Biol. 2016, 8, a005900.

[3] C. Manzo, M. F. Garcia-Parajo, Rep. Prog. Phys. 2015, 78, 124601.

[4] A. Kusumi, T. A. Tsunoyama, K. M. Hirosawa, R. S. Kasai, T. K. Fujiwara, Nat. Chem. Biol. 2014, 10, 524–532.

[5] J.-B. Sibarita, Histochem. Cell Biol. 2014, 141, 587–595.

[6] S. Wilmes, O. Beutel, Z. Li, V. Francois-Newton, C. P. Richter, D. Janning, C. Kroll, P. Hanhart, K. Hötte, C. You, G. Uzé, S. Pellegrini, J. Piehler, J. Cell Biol. 2015, 209, 579–593.

[7] J. Sotolongo Bellón, O. Birkholz, C. P. Richter, F. Eull, H. Kenneweg, S. Wilmes, U. Rothbauer, C. You, M. R. Walter, R. Kurre, J. Piehler, Cell Rep Methods 2022, 2, 100165.

[8] H. Shen, L. J. Tauzin, R. Baiyasi, W. Wang, N. Moringo, B. Shuang, C. F. Landes, Chem Rev. 2017, 117, 7331–7376.

[9] R. Iino, I. Koyama, A. Kusumi, Biophys. J. 2001, 80, 2667–2677.

[10] P. Winckler, L. Lartigue, G. Giannone, F. De Giorgi, F. Ichas, J.-B. Sibarita, B. Lounis, L. Cognet, Sci. Rep. 2013, 3, 2387.

[11] M.-L. I. E. Harwardt, P. Young, W. M. Bleymüller, T. Meyer, C. Karathanasis, H. H. Niemann, M. Heilemann, M. S. Dietz, FEBS Open Bio 2017, 7, 1422–1440.

[12] G. Giannone, E. Hosy, F. Levet, A. Constals, K. Schulze, A. I. Sobolevsky, M. P. Rosconi, E. Gouaux, R. Tampé, D. Choquet, L. Cognet, Biophys. J. 2010, 99, 1303–1310.

[13] C. Catapano, J. V. Rahm, M. Omer, L. Teodori, J. Kjems, M. S. Dietz, M. Heilemann, Cell. Mol. Life Sci. 2023, 80, 158.

[14] D. Albrecht, C. M. Winterflood, H. Ewers, Methods Appl Fluoresc 2015, 3, 024001.

[15] G. V. Los, L. P. Encell, M. G. McDougall, D. D. Hartzell, N. Karassina, C. Zimprich, M. G. Wood, R. Learish, R. F. Ohana, M. Urh, D. Simpson, J. Mendez, K. Zimmerman, P. Otto, G. Vidugiris, J. Zhu, A. Darzins, D. H. Klaubert, R. F. Bulleit, K. V. Wood, ACS Chem. Biol. 2008, 3, 373–382.

[16] R. F. Ohana, L. P. Encell, K. Zhao, D. Simpson, M. R. Slater, M. Urh, K. V. Wood, Protein Expr. Purif. 2009, 68, 110–120.

[17] V. Paakinaho, D. M. Presman, D. A. Ball, T. A. Johnson, R. L. Schiltz, P. Levitt, D. Mazza, T. Morisaki, T. S. Karpova, G. L. Hager, Nat. Commun. 2017, 8, 15896.

[18] A. Keppler, S. Gendreizig, T. Gronemeyer, H. Pick, H. Vogel, K. Johnsson, Nat. Biotechnol. 2003, 21, 86–89.

[19] P. J. Bosch, I. R. Corrêa Jr, M. H. Sonntag, J. Ibach, L. Brunsveld, J. S. Kanger, V. Subramaniam, Biophys. J. 2014, 107, 803–814.

[20] S. Manley, J. M. Gillette, G. H. Patterson, H. Shroff, H. F. Hess, E. Betzig, J. Lippincott-Schwartz, Nat. Methods 2008, 5, 155–157.

[21] Y. Yu, M. Li, Y. Yu, ACS Nano 2019, 13, 10860–10868.

[22] F. Stehr, J. Stein, J. Bauer, C. Niederauer, R. Jungmann, K. Ganzinger, P. Schwille, Nat. Commun. 2021, 12, 4432.

[23] C. Niederauer, C. Nguyen, M. Wang-Henderson, J. Stein, S. Strauss, A. Cumberworth, F. Stehr, R. Jungmann, P. Schwille, K. A. Ganzinger, Nat. Commun. 2023, 14, 4345.

[24] L. Albertazzi, M. Heilemann, Angew. Chem. Int. Ed Engl. 2023, 62, e202303390.

[25] M. Holtmannspötter, E. Wienbeuker, T. Dellmann, I. Watrinet, A. J. Garcia-Sáez, K. Johnsson, R. Kurre, J. Piehler, Angew. Chem. Int. Ed Engl. 2023, 62, e202219050.

[26] H. Hass, K. Masson, S. Wohlgemuth, V. Paragas, J. E. Allen, M. Sevecka, E. Pace, J. Timmer, J. Stelling, G. MacBeath, B. Schoeberl, A. Raue, NPJ Syst Biol Appl 2017, 3, 27.

[27] B. Blagoev, S.-E. Ong, I. Kratchmarova, M. Mann, Nat. Biotechnol. 2004, 22, 1139–1145.

[28] B. da Rocha-Azevedo, S. Lee, A. Dasgupta, A. R. Vega, L. R. de Oliveira, T. Kim, M. Kittisopikul, Z. A. Malik, K. Jaqaman, Cell Rep. 2020, 32, 108187.

[29] J. Kompa, J. Bruins, M. Glogger, J. Wilhelm, M. S. Frei, M. Tarnawski, E. D’Este, M. Heilemann, J. Hiblot, K. Johnsson, J. Am. Chem. Soc. 2023, 145, 3075–3083.

[30] S. Jang, K. K. Narayanasamy, J. V. Rahm, A. Saguy, J. Kompa, M. S. Dietz, K. Johnsson, Y. Shechtman, M. Heilemann, Biophys Rep (N Y) 2023, 3, 100123.

[31] M. Glogger, D. Wang, J. Kompa, A. Balakrishnan, J. Hiblot, H.-D. Barth, K. Johnsson, M. Heilemann, ACS Nano 2022, 16, 17991–17997.

[32] S. Bhatia, M. Edidin, S. C. Almo, S. G. Nathenson, Immunol. Lett. 2006, 104, 70–75.

[33] P. S. Linsley, S. G. Nadler, J. Bajorath, R. Peach, H. T. Leung, J. Rogers, J. Bradshaw, M. Stebbins, G. Leytze, W. Brady, A. R. Malacko, H. Marquardt, S.-Y. Shaw, J. Biol. Chem. 1995, 270, 15417–15424.

[34] J. Ibach, Y. Radon, M. Gelléri, M. H. Sonntag, L. Brunsveld, P. I. H. Bastiaens, P. J. Verveer, PLoS One 2015, 10, e0143162.

[35] M.-L. I. E. Harwardt, M. S. Schröder, Y. Li, S. Malkusch, P. Freund, S. Gupta, N. Janjic, S. Strauss, R. Jungmann, M. S. Dietz, M. Heilemann, Int. J. Mol. Sci. 2020, 21, DOI 10.3390/ijms21082803.

[36] Y. Yarden, M. X. Sliwkowski, Nat. Rev. Mol. Cell Biol. 2001, 2, 127–137.

[37] T. N. Baldering, C. Karathanasis, M.-L. I. E. Harwardt, P. Freund, M. Meurer, J. V. Rahm, M. Knop, M. S. Dietz, M. Heilemann, iScience 2021, 24, 101895.

[38] S. Wilmes, M. Hafer, J. Vuorio, J. A. Tucker, H. Winkelmann, S. Löchte, T. A. Stanly, K. D. Pulgar Prieto, C. Poojari, V. Sharma, C. P. Richter, R. Kurre, S. R. Hubbard, K. C. Garcia, I. Moraga, I. Vattulainen, I. S. Hitchcock, J. Piehler, Science 2020, 367, 643–652.

[39] K. C. Mudumbi, E. A. Burns, D. J. Schodt, Z. O. Petrova, A. Kiyatkin, L. W. Kim, E. M. Mangiacapre, I. Ortiz-Caraveo, H. Rivera Ortiz, C. Hu, K. D. Ashtekar, K. A. Lidke, D. S. Lidke, M. A. Lemmon, Cell Rep. 2024, 43, 113603.

[40] M. J. Saxton, Biophys. J. 1997, 72, 1744–1753.

[41] L. Weimann, K. A. Ganzinger, J. McColl, K. L. Irvine, S. J. Davis, N. J. Gay, C. E. Bryant, D. Klenerman, PLoS One 2013, 8, e64287.

[42] J. V. Rahm, S. Malkusch, U. Endesfelder, M. S. Dietz, M. Heilemann, Front. Comput. Sci. 2021, 3, DOI 10.3389/fcomp.2021.757653.

[43] M. Philippi, J. Dohle, I. Watrinet, M. Holtmannspötter, J. Li, O. Birkholz, Y. Miao, U. Rothbauer, K. Christopher Garcia, R. Kurre, J. Piehler, C. You, bioRxiv preprint 2024, DOI: 10.1101/2024.06.18.599024.

[44] D. T. Clarke, M. L. Martin-Fernandez, Methods Protoc 2019, 2, DOI 10.3390/mps2010012.

[45] C. Franco Nitta, E. W. Green, E. D. Jhamba, J. M. Keth, I. Ortiz-Caraveo, R. M. Grattan, D. J. Schodt, A. C. Gibson, A. Rajput, K. A. Lidke, B. S. Wilson, M. P. Steinkamp, D. S. Lidke, Elife 2021, 10, DOI 10.7554/eLife.63678.

[46] J. A. Rybak, A. R. Sahoo, S. Kim, R. J. Pyron, S. B. Pitts, S. Guleryuz, A. W. Smith, M. Buck, F. N. Barrera, J. Biol. Chem. 2023, 299, 104914.

[47] D.-H. Kim, S. Park, D.-K. Kim, M. G. Jeong, J. Noh, Y. Kwon, K. Zhou, N. K. Lee, S. H. Ryu, PLoS Biol. 2018, 16, e2006660.

[48] Y. Huang, S. Bharill, D. Karandur, S. M. Peterson, M. Marita, X. Shi, M. J. Kaliszewski, A. W. Smith, E. Y. Isacoff, J. Kuriyan, Elife 2016, 5, DOI: 10.7554/eLife.14107.

[49] S. T. Low-Nam, K. A. Lidke, P. J. Cutler, R. C. Roovers, P. M. P. van Bergen en Henegouwen, B. S. Wilson, D. S. Lidke, Nat. Struct. Mol. Biol. 2011, 18, 1244–1249.

[50] G. Orr, D. Hu, S. Ozçelik, L. K. Opresko, H. S. Wiley, S. D. Colson, Biophys. J. 2005, 89, 1362–1373.

[51] A. I. König, R. Sorkin, A. Alon, D. Nachmias, K. Dhara, G. Brand, O. Yifrach, E. Arbely, Y. Roichman, N. Elia, Nanoscale 2020, 12, 3236–3248.

[52] P. Nagy, J. Claus, T. M. Jovin, D. J. Arndt-Jovin, Proc. Natl. Acad. Sci. U. S. A. 2010, 107, 16524–16529.

[53] X. Bai, P. Sun, X. Wang, C. Long, S. Liao, S. Dang, S. Zhuang, Y. Du, X. Zhang, N. Li, K. He, Z. Zhang, Cell Discov 2023, 9, 18.

[54] J. Icha, M. Weber, J. C. Waters, C. Norden, Bioessays 2017, 39, DOI 10.1002/bies.201700003.

[55] M. Weigert, U. Schmidt, T. Boothe, A. Müller, A. Dibrov, A. Jain, B. Wilhelm, D. Schmidt, C. Broaddus, S. Culley, M. Rocha-Martins, F. Segovia-Miranda, C. Norden, R. Henriques, M. Zerial, M. Solimena, J. Rink, P. Tomancak, L. Royer, F. Jug, E. W. Myers, Nat. Methods 2018, 15, 1090–1097.

[56] P. Kefer, F. Iqbal, M. Locatelli, J. Lawrimore, M. Zhang, K. Bloom, K. Bonin, P.-A. Vidi, J. Liu, Mol. Biol. Cell 2021, 32, 903–914.

[57] V. Glembockyte, J. Lin, G. Cosa, J. Phys. Chem. B 2016, 120, 11923–11929.

[58] V. Glembockyte, R. Wieneke, K. Gatterdam, Y. Gidi, R. Tampé, G. Cosa, J. Am. Chem. Soc. 2018, 140, 11006–11012.

[59] M. Isselstein, L. Zhang, V. Glembockyte, O. Brix, G. Cosa, P. Tinnefeld, T. Cordes, J. Phys. Chem. Lett. 2020, 11, 4462–4480.

[60] R. B. Altman, D. S. Terry, Z. Zhou, Q. Zheng, P. Geggier, R. A. Kolster, Y. Zhao, J. A. Javitch, J. D. Warren, S. C. Blanchard, Nat. Methods 2011, 9, 68–71.

[61] P. Tinnefeld, T. Cordes, Nat. Methods 2012, 9, 426–7; author reply 427–8.

[62] A. Sergé, N. Bertaux, H. Rigneault, D. Marguet, Nat. Methods 2008, 5, 687–694.

[63] M. S. Frei, P. Hoess, M. Lampe, B. Nijmeijer, M. Kueblbeck, J. Ellenberg, H. Wadepohl, J. Ries, S. Pitsch, L. Reymond, K. Johnsson, Nat. Commun. 2019, 10, 4580.

[64] J. Kompa, J. Bruins, M. Glogger, J. Wilhelm, M. S. Frei, M. Tarnawski, E. D’Este, M. Heilemann, J. Hiblot, K. Johnsson, J. Am. Chem. Soc. 2023, 145, 3075–3083.

[65] G. Lukinavičius, K. Umezawa, N. Olivier, A. Honigmann, G. Yang, T. Plass, V. Mueller, L. Reymond, I. R. Corrêa Jr, Z.-G. Luo, C. Schultz, E. A. Lemke, P. Heppenstall, C. Eggeling, S. Manley, K. Johnsson, Nat. Chem. 2013, 5, 132–139.

[66] F. Fricke, J. Beaudouin, R. Eils, M. Heilemann, Sci. Rep. 2015, 5, 14072.

[67] R. E. Carter, A. Sorkin, J. Biol. Chem. 1998, 273, 35000–35007.

[68] D. G. Gibson, L. Young, R.-Y. Chuang, J. C. Venter, C. A. Hutchison 3rd, H. O. Smith, Nat. Methods 2009, 6, 343–345.

[69] M. J. Malecki, C. Sanchez-Irizarry, J. L. Mitchell, G. Histen, M. L. Xu, J. C. Aster, S. C. Blacklow, Mol. Cell. Biol. 2006, 26, 4642–4651.

[70] A. Edelstein, N. Amodaj, K. Hoover, R. Vale, N. Stuurman, Curr. Protoc. Mol. Biol. 2010, Chapter 14, Unit14.20.

[71] M. Ovesný, P. Křížek, J. Borkovec, Z. Svindrych, G. M. Hagen, Bioinformatics 2014, 30, 2389–2390.

[72] J. Schindelin, I. Arganda-Carreras, E. Frise, V. Kaynig, M. Longair, T. Pietzsch, S. Preibisch, C. Rueden, S. Saalfeld, B. Schmid, J.-Y. Tinevez, D. J. White, V. Hartenstein, K. Eliceiri, P. Tomancak, A. Cardona, Nat. Methods 2012, 9, 676–682.

[73] M. Endesfelder, C. Schießl, B. Turkowyd, T. Lechner, U. Endesfelder, manuscript in preparation.

