## Supplementary Information for "Long-term single-molecule tracking in living cells using weak-affinity protein labeling"

### Supporting Figures

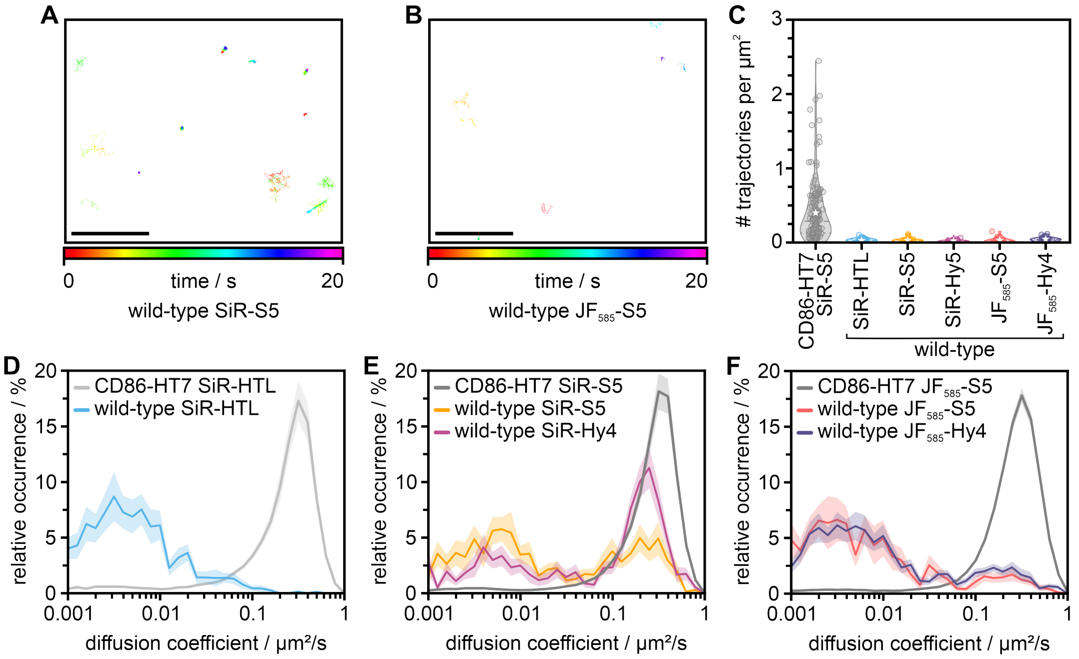


**Figure S1:** U-2 OS wild-type cells imaged with covalent and exchangeable HTLs as negative controls. Single-molecule trajectories of exemplary cells acquired using the (A) SiR-S5 and (B) JF585-S5. Trajectories are color-coded over the 20 s acquisition time. Scale bar 5 µm.

(C) Number of trajectories per area for each ligand in U-2 OS wild-type cells compared to CD86-HT7 imaged with SiR-S5. Very few trajectories were detected in the negative controls. Dashed lines represent the median, stars the mean, and dotted lines the interquartile range.

(D-F) Relative occurrence of the mean diffusion coefficient per cell for each ligand in U-2 OS wild-type cells compared to the respective CD86-HT7 cell line.

All errors represent the SEM (N = 20 cells for negative controls, N = 160 cells for CD86-HT7 imaged with SiR-S5 and SiR-HTL, N = 180 cells for CD86-HT7 imaged with JF585-S5).

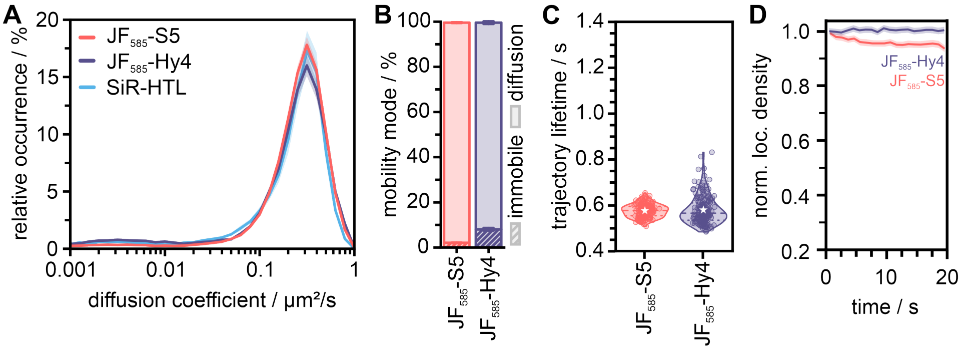


**Figure S2:** CD86-HT7 and CD86-dHT7 imaged with the xHTLs JF585-S5 and JF585-Hy4.

(A) Relative occurrence of the mean diffusion coefficient per cell for JF585-tagged exchangeable HaloTag Ligands.

(B) Percentage of mobility modes per cell for JF585-S5 and JF585-Hy4. Single-molecule trajectories were assigned to the classes immobile or diffusion.

(C) Lifetime of single-molecule trajectories of the non-covalent JF585-S5 binding to the HaloTag7 and JF585-Hy4 binding to the dead mutant of the HaloTag7. Dashed lines represent the median, stars the mean, and dotted lines the interquartile range.

(D) Mean number of localizations per area binned into 1 s intervals, normalized to the respective data in the first frame and plotted over time for JF585-S5 (red) and JF585-Hy4 (purple).

All errors represent the SEM (N = 180 cells for all conditions).

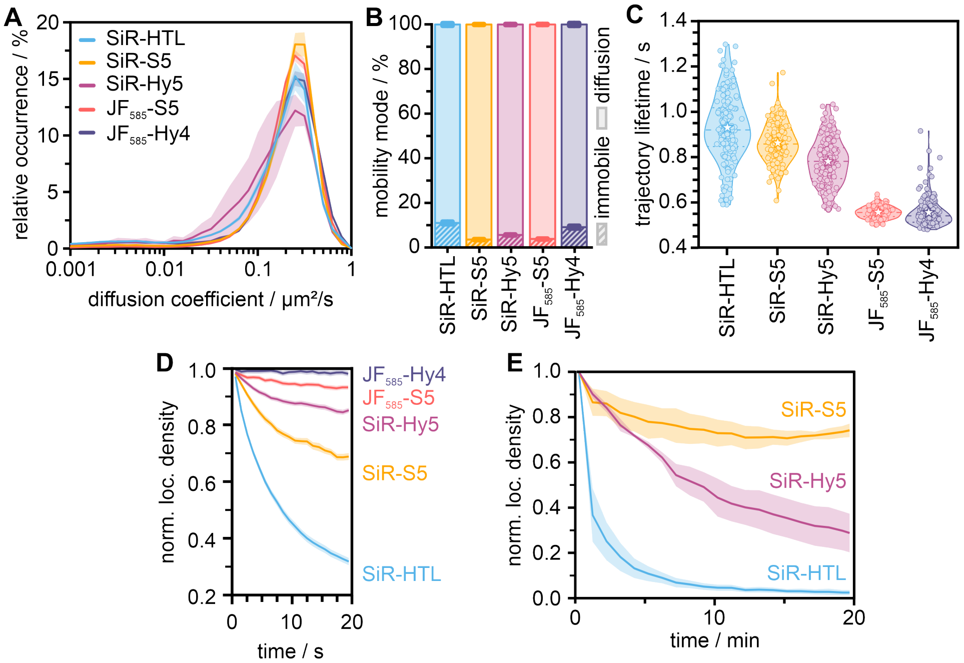


**Figure S3:** CTLA-4-HT7 and CTLA-4-dHT7 imaged with the covalent HaloTag Ligand SiR-HTL as well as the xHTLs SiR-S5, SiR-Hy5, $JF_{585}$-S5, and $JF_{585}$-Hy4.

(A) Relative occurrence of the mean diffusion coefficient per cell for SiR-HTL (blue), SiR-S5 (yellow), SiR-Hy5 (magenta), JF585-S5 (red), and JF585-Hy4 (purple).

(B) Percentage of mobility modes per cell for the different ligands. Single-molecule trajectories were assigned to the classes immobile or diffusion.

(C) Lifetime of single-molecule trajectories of the different ligands. Dashed lines represent the median, stars the mean, and dotted lines the interquartile range.

(D) Mean number of localizations per area from short SPT measurements binned into 1 s intervals, normalized to the respective data in the first frame, and plotted over time for the different ligands.

(E) Mean number of localizations per area from long-time SPT measurements binned into 1 min intervals, normalized to the respective data in the first frame, and plotted over time for SiR-HTL (blue), SiR-S5 (yellow), and SiR-Hy5 (magenta).

All errors represent the SEM (N = 160 cells for SiR-tagged ligands and N = 180 cells for $JF_{585}$-tagged ligands in A-D, and N = 4 cells for E)

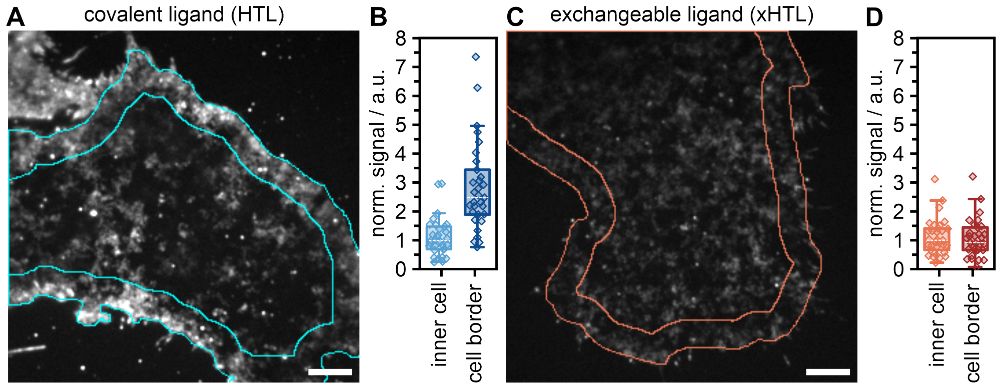


**Figure S4:** Comparison of the fluorescence signal in cells imaged with the covalent ligand SiR-HTL (A, B) and the exchangeable probe SiR-S5 (C, D). For the analysis, a full 20 s SPT measurement was z-projected with the setting "standard deviation" in Fiji (A, C) and the median intensity value measured either in the region of the border or inside of the cell (B, D). The median intensity values per cell were normalized to the overall median intensity inside the cells per condition. 31 cells were analyzed for each condition. The box plots display the median (dashed white line) with the 25th and 75th percentile with whiskers reaching to the last data point within the 1.5× interquartile range, and diamonds represent single-cell values.

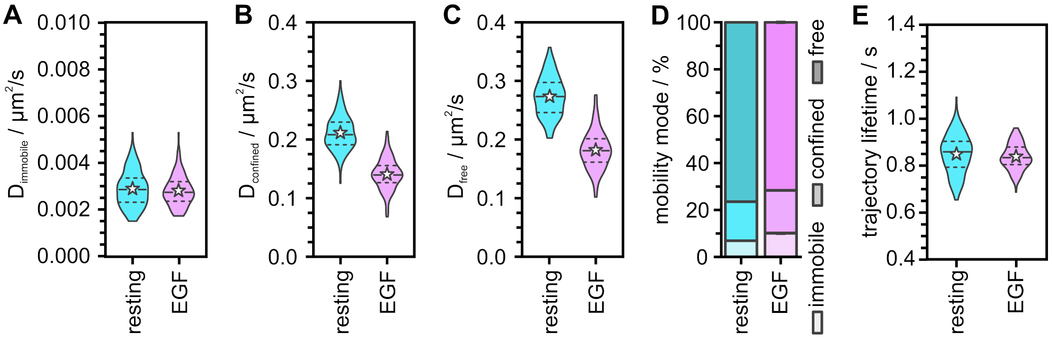


**Figure S5:** Diffusion properties of EGFR-HT7 in resting and EGF-stimulated U-2 OS cells. SiR-S5 was used as xHTL.

(A-C) Distribution of diffusion coefficients for the individual diffusion modes immobile (A), confined (B), and free (C). Dashed lines represent the median, dotted lines represent the quartiles, and stars represent the mean. In the data seen in **Figure 3**, the confined and freely moving receptor fractions were pooled.

(D) Occurrence of immobile molecules and confined and freely diffusing EGFR-HT7.

(E) Lifetime of single-molecule trajectories extracted from SPT data of EGFR-HT7. Dashed lines represent the median, stars the mean, and dotted lines the interquartile range.

160 cells were analyzed per condition for all plots.

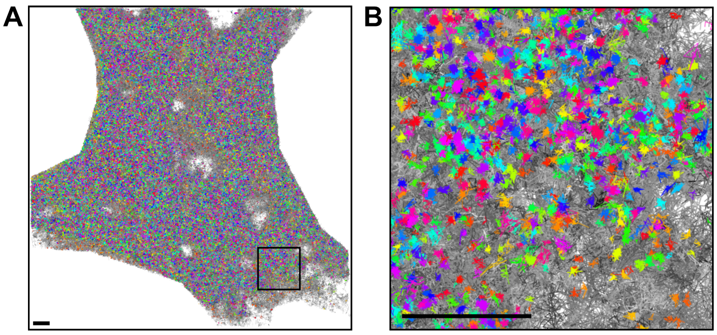


**Figure S6:** Related to Figure 3FG. (A) Overlay of all trajectories (gray) and only immobile trajectories (rainbow, randomly color-coded) throughout a 30 min measurement of EGFR-HT7. (B) Zoom-in of the region marked in (A). Scale bars 2 µm.

### Supporting Tables

**Supplementary Table S1:** Mean diffusion coefficients of all measured protein-ligand combinations calculated from 20 s measurements. The diffusion coefficient of mobile particles ($D_{mobile}$) is calculated as the mean value of the diffusion coefficient of the confined ($D_{confined}$) and free ($D_{free}$) mobilities modes weighted by their relative occurrence (**Table S2**). The global diffusion coefficient $D_{global}$ is calculated similarly including all three mobility types (immobile, confined, free). POI: protein of interest. All given errors are the SEM.

| POI | ligand | $D_{global}$ / µm²s⁻¹ | $D_{immobile}$ / µm²s⁻¹ | $D_{mobile}$ / µm²s⁻¹ | $D_{confined}$ / µm²s⁻¹ | $D_{free}$ / µm²s⁻¹ |
|---|---|---|---|---|---|---|
| CD86-HT7 | SiR-HTL | 0.294 ± 0.007 | 0.0043 ± 0.0009 | 0.315 ± 0.007 | 0.245 ± 0.016 | 0.336 ± 0.007 |
| CD86-HT7 | SiR-S5 | 0.300 ± 0.011 | 0.0036 ± 0.0010 | 0.319 ± 0.010 | 0.262 ± 0.03 | 0.331 ± 0.011 |
| CD86-HT7 | $JF_{585}$-S5 | 0.333 ± 0.008 | 0.0046 ± 0.0013 | 0.344 ± 0.008 | 0.296 ± 0.016 | 0.360 ± 0.009 |
| CD86-dHT7 | SiR-Hy5 | 0.283 ± 0.005 | 0.0034 ± 0.0006 | 0.300 ± 0.005 | 0.245 ± 0.013 | 0.312 ± 0.006 |
| CD86-dHT7 | $JF_{585}$-Hy4 | 0.31 ± 0.02 | 0.0044 ± 0.0017 | 0.34 ± 0.02 | 0.29 ± 0.04 | 0.36 ± 0.02 |
| CTLA-4-HT7 | SiR-HTL | 0.228 ± 0.008 | 0.0051 ± 0.0011 | 0.255 ± 0.008 | 0.194 ± 0.017 | 0.274 ± 0.009 |
| CTLA-4-HT7 | SiR-S5 | 0.270 ± 0.005 | 0.0042 ± 0.0011 | 0.279 ± 0.005 | 0.235± 0.012 | 0.288 ± 0.005 |
| CTLA-4-HT7 | $JF_{585}$-S5 | 0.285 ± 0.005 | 0.007 ± 0.002 | 0.291 ± 0.005 | 0.249 ± 0.009 | 0.303 ± 0.006 |
| CTLA-4-dHT7 | SiR-Hy5 | 0.257 ± 0.004 | 0.0041 ± 0.0006 | 0.271 ± 0.004 | 0.222 ± 0.009 | 0.283 ± 0.004 |
| CTLA-4-dHT7 | $JF_{585}$-Hy4 | 0.272 ± 0.015 | 0.006 ± 0.002 | 0.295 ± 0.016 | 0.248 ± 0.03 | 0.311 ± 0.018 |
| EGFR-HT7 rest. | SiR-S5 | 0.245 ± 0.004 | 0.0029 ± 0.0005 | 0.262 ± 0.004 | 0.211 ± 0.010 | 0.273 ± 0.004 |
| EGFR-HT7 +EGF | SiR-S5 | 0.158 ± 0.003 | 0.0028 ± 0.0004 | 0.174 ± 0.003 | 0.140 ± 0.007 | 0.182 ± 0.004 |

**Supplementary Table S2:** Mean relative occurrence of mobility modes of all measured protein-ligand combinations calculated from 20 s measurements. The mobile fraction is the sum of the confined and free mobility modes. All given errors are the SEM.

| POI | ligand | immobile / % | mobile / % | confined / % | free / % |
|---|---|---|---|---|---|
| CD86-HT7 | SiR-HTL | 7.6 ± 0.3 | 92.4 ± 0.3 | 18.4 ± 0.3 | 74.0 ± 0.4 |
| CD86-HT7 | SiR-S5 | 6.1 ± 0.3 | 93.4 ± 0.4 | 16.8 ± 0.2 | 77.1 ± 0.3 |
| CD86-HT7 | $JF_{585}$-S5 | 3.6 ± 0.2 | 98.07 ± 0.15 | 22.5 ± 0.2 | 73.9 ± 0.3 |
| CD86-dHT7 | SiR-Hy5 | 6.6 ± 0.4 | 93.9 ± 0.3 | 16.0 ± 0.3 | 77.4 ± 0.4 |
| CD86-dHT7 | $JF_{585}$-Hy4 | 9.0 ± 0.6 | 92.0 ± 0.6 | 22.5 ± 0.4 | 68.5 ± 0.7 |
| CTLA-4-HT7 | SiR-HTL | 10.8 ± 0.6 | 89.2 ± 0.6 | 20.5 ± 0.3 | 68.6 ± 0.7 |
| CTLA-4-HT7 | SiR-S5 | 3.4 ± 0.2 | 96.6 ± 0.2 | 16.9 ± 0.2 | 79.8 ± 0.3 |
| CTLA-4-HT7 | $JF_{585}$-S5 | 1.9 ± 0.2 | 96.4 ± 0.2 | 23.9 ± 0.2 | 74.1 ± 0.2 |
| CTLA-4-dHT7 | SiR-Hy5 | 5.4 ± 0.2 | 94.6 ± 0.2 | 18.9 ± 0.2 | 75.7 ± 0.3 |
| CTLA-4-dHT7 | $JF_{585}$-Hy4 | 8.0 ± 0.6 | 91.0 ± 0.6 | 23.9 ± 0.3 | 68.1 ± 0.6 |
| EGFR-HT7 rest. | SiR-S5 | 6.9 ± 0.2 | 93.1 ± 0.2 | 16.8 ± 0.2 | 76.3 ± 0.2 |
| EGFR-HT7 ±EGF | SiR-S5 | 10.1 ± 0.4 | 89.9 ± 0.4 | 17.50 ± 0.14 | 72.44 ± 0.4 |

**Supplementary Table S3:** Mean trajectory lifetime of all measured protein-ligand combinations given in seconds calculated from 20 s measurements. The mean value of the global lifetime is calculated as the mean value of the values extracted from trajectories classified as immobile, confined, and freely diffusing weighted by their relative occurrence (**Table S2**). All given errors are the SEM.

| POI | ligand | trajectory life-time global / s | trajectory life-time immobile / s | trajectory life-time confined / s | trajectory life-time free / s |
|---|---|---|---|---|---|
| CD86-HT7 | SiR-HTL | 0.94 ± 0.03 | 0.88 ± 0.10 | 0.63 ± 0.03 | 1.02 ± 0.04 |
| CD86-HT7 | SiR-S5 | 0.93 ± 0.04 | 0.85 ± 0.16 | 0.64 ± 0.04 | 0.99 ± 0.05 |
| CD86-HT7 | $JF_{585}$-S5 | 0.576 ± 0.010 | 0.68 ± 0.10 | 0.522 ± 0.013 | 0.586 ± 0.012 |
| CD86-dHT7 | SiR-Hy5 | 0.87 ± 0.02 | 0.83 ± 0.09 | 0.62 ± 0.02 | 0.92 ± 0.02 |
| CD86-dHT7 | $JF_{585}$-Hy4 | 0.58 ± 0.03 | 0.76 ± 0.18 | 0.51 ± 0.03 | 0.57 ± 0.03 |
| CTLA-4-HT7 | SiR-HTL | 0.93 ± 0.04 | 0.89 ± 0.12 | 0.62 ± 0.03 | 1.02 ± 0.05 |
| CTLA-4-HT7 | SiR-S5 | 0.86 ± 0.02 | 0.75 ± 0.09 | 0.62 ± 0.02 | 0.91 ± 0.02 |
| CTLA-4-HT7 | $JF_{585}$-S5 | 0.556 ± 0.006 | 0.60 ± 0.07 | 0.513 ± 0.009 | 0.567 ± 0.008 |
| CTLA-4-dHT7 | SiR-Hy5 | 0.780 ± 0.012 | 0.74 ± 0.05 | 0.600 ± 0.014 | 0.825 ± 0.015 |
| CTLA-4-dHT7 | $JF_{585}$-Hy4 | 0.56 ± 0.02 | 0.71 ± 0.13 | 0.50 ± 0.02 | 0.55 ± 0.03 |
| EGFR-HT7 rest. | SiR-S5 | 0.85 ± 0.02 | 0.76 ± 0.05 | 0.63 ± 0.02 | 0.90 ± 0.02 |
| EGFR-HT7 ±EGF | SiR-S5 | 0.85 ± 0.02 | 0.75 ± 0.04 | 0.63 ± 0.02 | 0.90 ± 0.02 |
